# The Almond plumage color pattern is associated with eye pigmentation defects in the domestic pigeon (*Columba livia*)

**DOI:** 10.1101/2020.10.17.339655

**Authors:** Rebecca Bruders, Max Sidesinger, Michael D. Shapiro

**Affiliations:** School of Biological Sciences, University of Utah, Salt Lake City, Utah, United States of America

## Abstract

Changes in epidermal pigmentation are associated with eye defects in humans and other vertebrates. In the rock pigeon (*Columba livia*), the sex-linked Almond color pattern is characterized by hypopigmentation of epidermal structures. The trait is controlled by the classical *Stipper* (*St*) locus, and homozygous (*Z^St^Z^St^*) Almond males often have severe eye defects. Heterozygous (*Z^St^*Z^+^) and hemizygous (Z^*St*^W) pigeons do not typically have obvious eye defects, suggesting that higher dosage of the mutant allele is deleterious. Because Almond pigeons have pronounced hypopigmentation in epidermal structures, we hypothesized that they might also have reduced eye pigmentation. Here, we examined pigmentation in the iris, ciliary body, anterior retinal pigmented epithelium (RPE), and posterior RPE in pigeons with and without Almond alleles. We found that pigmentation of anterior segment structures was reduced in birds with at least one Almond allele. However, posterior eye pigmentation was substantially reduced only in homozygous Almond birds. We postulate that the gradient of effects on eye pigmentation is due to the different embryological origins of anterior and posterior eye pigment-producing cells.

## Introduction

Eye defects are linked with changes in epidermal pigmentation in a variety of vertebrate species [1–5]. This link is driven in part by shared mechanisms that regulate development of pigment in both the eye and epidermis, including appendages such as feathers, scales, and hairs. Melanin-producing cells synthesize pigment granules in specialized organelles called melanosomes, which are exported to cells in the eye and skin [6, 7]. As a consequence of the common organellar origins of eye and skin melanins, mutations in genes important for melanosome function can lead to melanin-based pigmentation defects in both eyes and epidermal derivatives. For example, in humans, mutations in several genes involved in pigmentation are associated with oculocutaneous albinism (OCA), a group of inherited diseases that affect eye, skin, and hair pigmentation [5, 8–12], In mice and zebrafish, mutations in melanosome genes such as *Slc24a5, Rab38*, and *Oca2* can result in pigmentation changes in both the eye and epidermal tissues that are phenotypically similar to OCA in humans [1, 13–15].

In addition to OCA-like syndromes, other mutations have linked effects on pigmentation of the eye, epidermis, and epidermal appendages. Dogs that are heterozygous for a mutation in *pmel* (a gene that encodes a melanosome structural protein) have a patchwork Merle coat color pattern. However, homozygous Merle dogs have a dramatic reduction in coat pigmentation and show variable eye defects, including microphthalmia, iris coloboma, and cataracts [16, 17].

Pigment development in the eye and epidermis has both shared and distinct characteristics, which probably explains the spectrum of shared and distinct effects of mutations on eye and epidermal phenotypes. Most epidermal pigmentation comes from melanin-producing cells called melanocytes. Melanocytes are neural crest-derived cells that produce pigment-loaded melanosomes [18, 19], which are subsequently transferred to the skin and its appendages [20]. In contrast, eye pigmentation originates from two distinct cell populations, melanocytes and pigmented epithelial cells. The latter population originates from optic cup neuroepithelium instead of neural crest [7, 19–21].

In the eye, melanosome-containing structures include the iris pigmented epithelium (IPE), ciliary body, and retinal pigment epithelium (RPE) (Fig 1). Pigmentation is protective against cellular oxidative damage due to UV radiation, yet pigmentation is not directly necessary for normal functions of eye structures [22–24]. The anterior structures of the eye (ciliary body and IPE) receive melanosomes that come from both the neural crest and optic cup neuroepithelium [7, 19–21]. However, the RPE of the posterior compartment of the eye receives pigmented melanosomes from only the optic cup neuroepithelium. While both types of cells produce melanosomes, neural crest cell-derived melanocytes and optic cup neuroepithelium-derived RPE cells sometimes respond differently to mutations in pigmentation genes. For example, in Dominant white chickens, mutations in *PMEL17* (*Pmel* ortholog) result in a lack of pigmentation in feathers and choroidal melanocytes (derived from neural crest cells), but RPE pigmentation is unaffected [25]. Additionally, mutations in the transcription factor gene *Mitf*, a “master regulator” of pigment development, can cause a lack of pigment in neural crest or optic cup derivatives – but not necessarily both – due to cell-type specific isoforms of Mitf [7, 26]. Finally, more than 150 genes are involved in epidermis and epidermal appendage pigmentation in laboratory mice, and some of these mutations affect eye development, while others do not [27]. The partially overlapping developmental origins of epidermal and eye pigmentation provide a general explanation for why some mutations might only lead to skin changes, while others induce differences in both organs.

**Figure 1.**
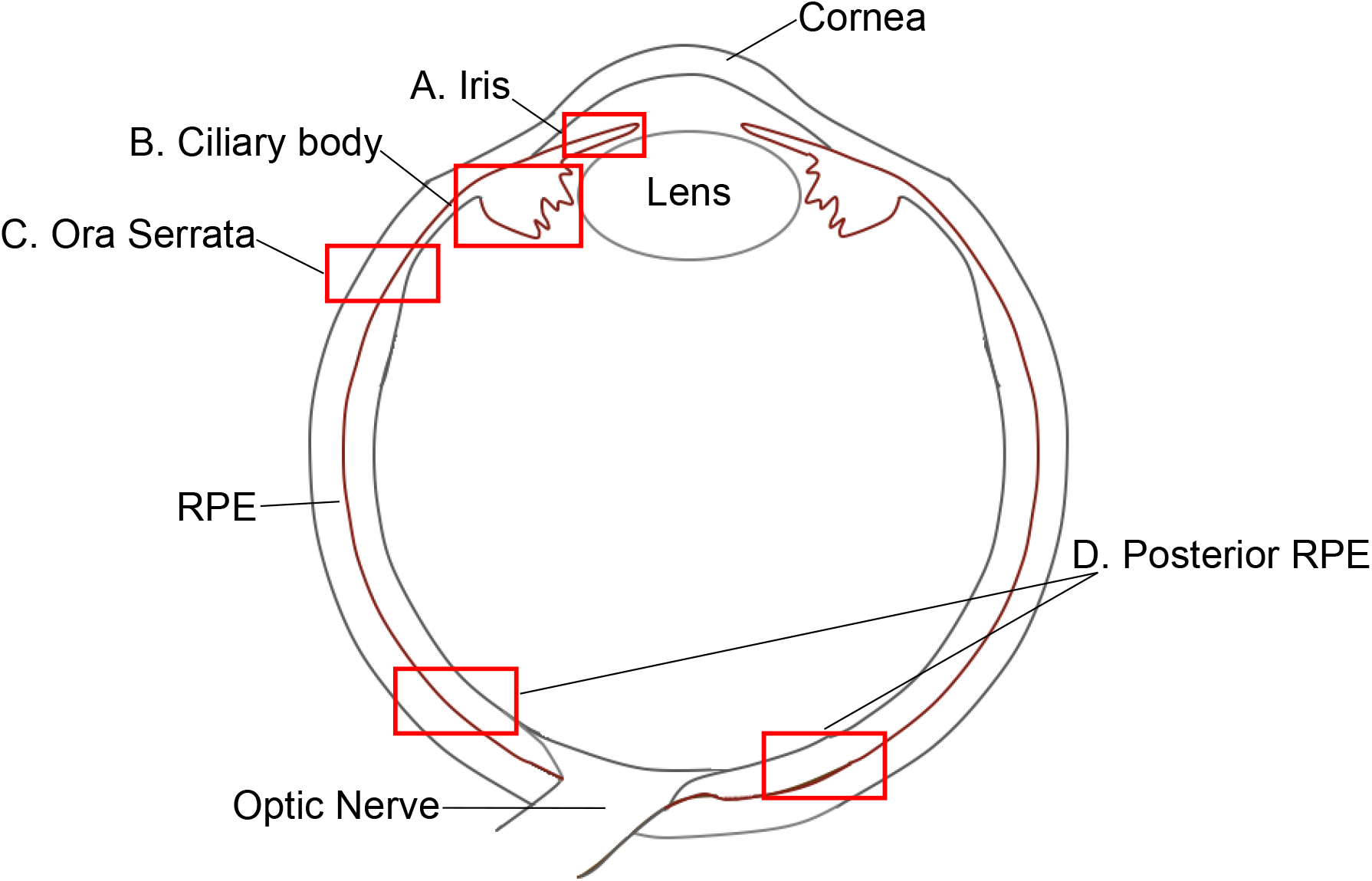
Simplified schematic of an eye in transverse section. Anterior is at the top, posterior at the bottom. Each region of the eye that was analyzed for pigmentation is labeled as follows: (A), Iris; (B), Ciliary body; (C), Ora serrata; (D), Posterior RPE. Red boxes represent approximate locations of regions that were imaged and analyzed (not to scale). Brown lines represent pigmented cell layers of the eye. RPE, retinal pigment epithelium. Schematic based on [72, 73].

In the domestic pigeon, Almond feather pattern is a variegated pattern of hypopigmented and heavily pigmented feathers that is also linked with eye defects. Almond is a sex-linked trait controlled by the *St* locus, in which hemizygous Almond females (ZW sex chromosomes) and heterozygous Almond males (ZZ) have changes in feather pigmentation that create a sprinkled pattern, but do not have known eye defects. However, homozygous Almond males typically have severe feather hypopigmentation and eye defects [28–30] described as “bulging bladder eyes” or “pop-eyes” [31–35] with “dark and irregular pupils” (coloboma) [34] or lens-absent eyes [32].

Previously, we identified a 359-kb candidate region for Almond on the Z-chromosome [36]. This region contains eight annotated protein-coding genes, none of which have fixed coding changes in Almond birds. However, we found that Almond genomes have a complex copy number variant in the candidate region that contains two full-length genes, *Mlana* and *Slc16a13*. Both genes have expression changes in the regenerating feather buds of Almond compared to non-Almond birds. Mlana is involved in melanosome development, and it interacts with Pmel to form complexes in the endoplasmic reticulum, trans-Golgi network, and stage I melanosomes [37, 38]. Mutations in *Pmel* are linked to eye, epidermis, and epidermal appendage pigmentation defects in numerous animals [16, 17, 25, 39–49]. In contrast, *Slc16a13* does not have a known role in pigmentation, and the function of this gene is poorly understood beyond its genomic association with type 2 diabetes [50–55]. We therefore proposed *Mlana* as a strong candidate gene for Almond [36].

Changes to feather pigmentation in Almond pigeons are well described, but whether plumage variation is linked to eye pigmentation changes remains unknown. Therefore, we performed a histologic study of eyes of pigeons with and without Almond alleles. Specifically, we quantified pigment content in the RPE, ciliary body, and iris to test for eye pigmentation changes associated with Almond genotypes.

## Results

### Iris

All eyes from pigeons with Almond genotypes had less pigmentation than non-Almonds in the iris, the most anterior structure we measured (Fig 1A and 2A; Anova, p = 2.2 x 10^-7^; post-hoc Tukey, p < 0.05 for all Almond genotypes). Homozygous Almonds had a mean of 1.0% of the iris pigmented, compared to 51.2% in birds without Almond alleles (post-hoc Tukey, p = 3.0 x 10^-7^). Heterozygotes and hemizygotes were more pigmented than homozygotes (9.2% and 9.7%, respectively), but the differences among genotypes were not significant (post-hoc Tukey, p > 0.05).

**Figure 2.**
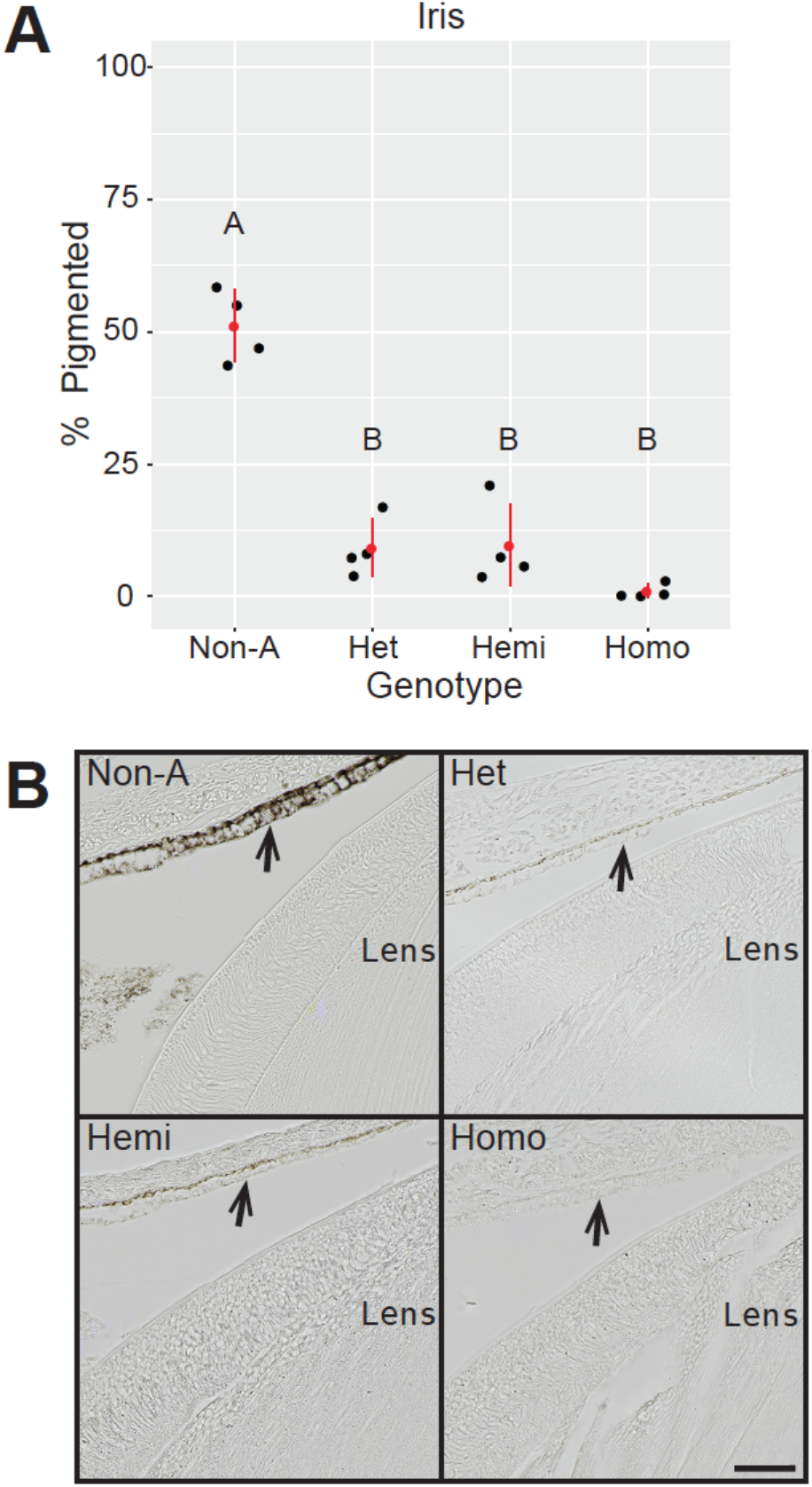
Iris pigmentation is reduced in all Almond genotypes. (A) Percentage of the IPE that is pigmented in different genotypes. Four hatchling eyes were analyzed per genotype. 3 sections were analyzed for each eye, and mean values are plotted as black dots. Red dot and line show mean and standard deviation, respectively. Different letters indicate groups with statistically significant differences in gene expression determined by Anova and post-hoc Tukey (p < 0.05). Genotypes: Non-A, non-Almond; Het, heterozygous Almond; Hemi, hemizygous Almond; Homo, homozygous Almond. (B) Representative images of the IPE for each genotype. Arrows point to the pigmented layer of the IPE. Pigmentation is reduced in heterozygous and hemizygous Almond hatchlings and is very faint in homozygous Almond hatchling eyes. Scale bar in bottom right image is 50 μm.

### Ciliary body

In the ciliary body, all Almond genotypes were less pigmented than non-Almonds (Fig 3A and 3B; Anova, p = 1.5 x 10^-8^, post-hoc Tukey, p > 0.05). Similar to the iris, the ciliary body had the highest mean pigmented area in non-Almond eyes (33.1%), and the lowest in homozygous Almond eyes (2.4%, Fig 3A; post-hoc Tukey, p = 0). Hemizygotes were slightly more pigmented than homozygotes with a mean of 6.4% pigmentation (post-hoc Tukey, p > 0.05). Heterozygotes had the most pigmentation of the Almond genotypes (9.7%, significantly higher than homozygous Almond; post-hoc Tukey, p = 0.019).

**Figure 3.**
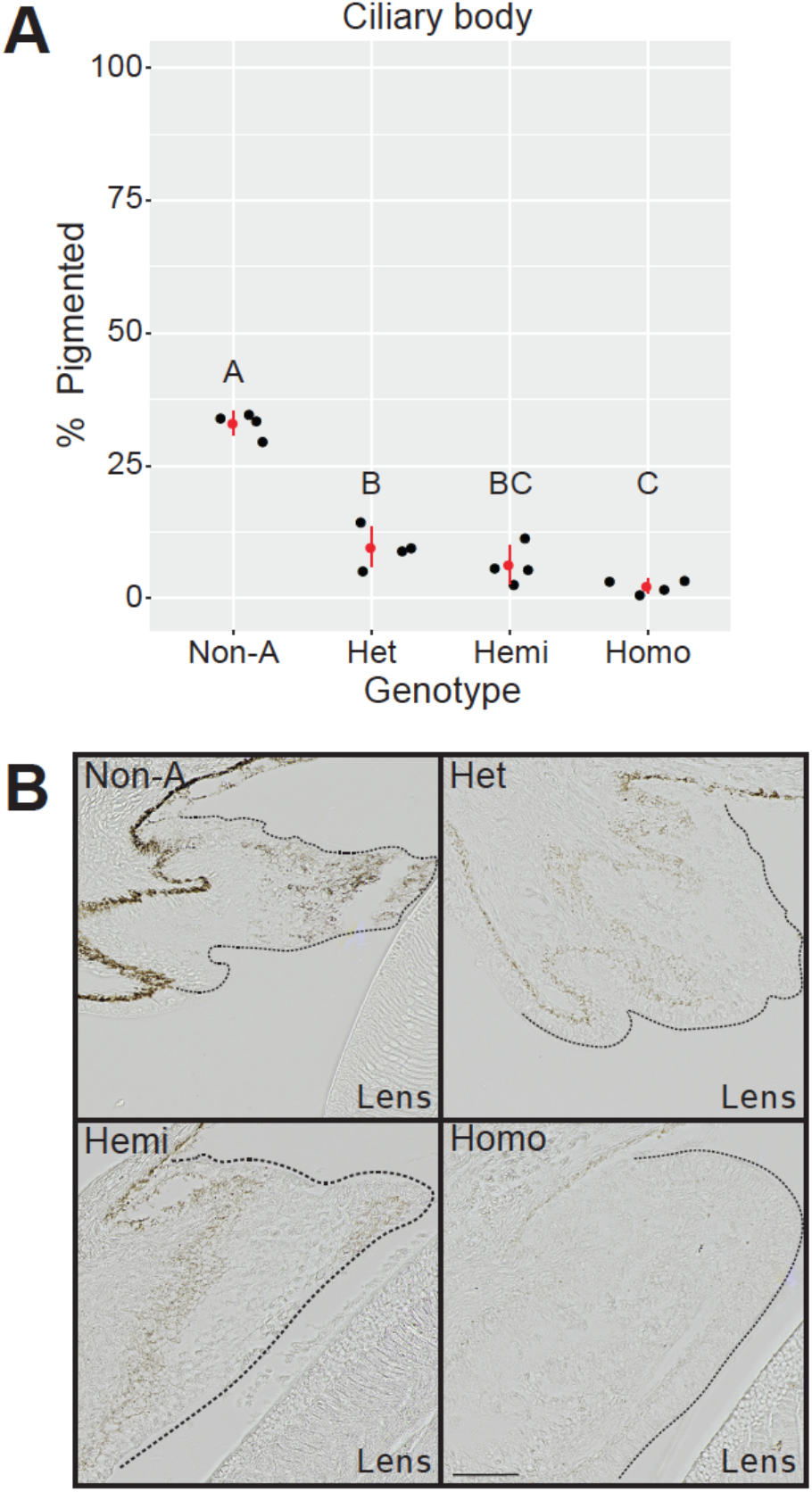
Ciliary body pigmentation is reduced in all Almond genotypes. (A) Percentage of the ciliary body that is pigmented in different genotypes. Four hatchling eyes were analyzed per genotype. 3 sections were analyzed for each eye, and mean values are plotted as black dots. Red dot and line show mean and standard deviation, respectively. Different letters indicate groups with statistically significant differences in gene expression determined by Anova and post-hoc Tukey (p < 0.05). Genotypes: Non-A, non-Almond; Het, heterozygous Almond; Hemi, hemizygous Almond; Homo, homozygous Almond. (B) Representative images of the ciliary body (outlined with dotted line) for each genotype. Pigmentation is reduced in heterozygous and hemizygous Almond hatchlings and is very faint in homozygous Almond hatchling eyes. Scale bar in bottom right image is 50 μm.

### Anterior RPE

Anterior RPE pigmentation was measured at the ora serrata, a landmark that can be reliably identified because of its distinct morphology (Fig 1C). The anterior RPE of all Almond genotypes had significantly less pigmented area compared to non-Almonds (Fig 4A and 4B; Anova, p = 1.4 x 10^-4^; post-hoc Tukey, p > 0.05). Non-Almond eyes had a mean of 51.9% pigmentation in the anterior RPE, while heterozygous, hemizygous, and homozygous eyes had means of 28.1%, 19.2%, and 5.7%, respectively. The anterior RPE was significantly less pigmented in homozygotes than in heterozygotes (post-hoc Tukey, p = 0.03).

**Figure 4.**
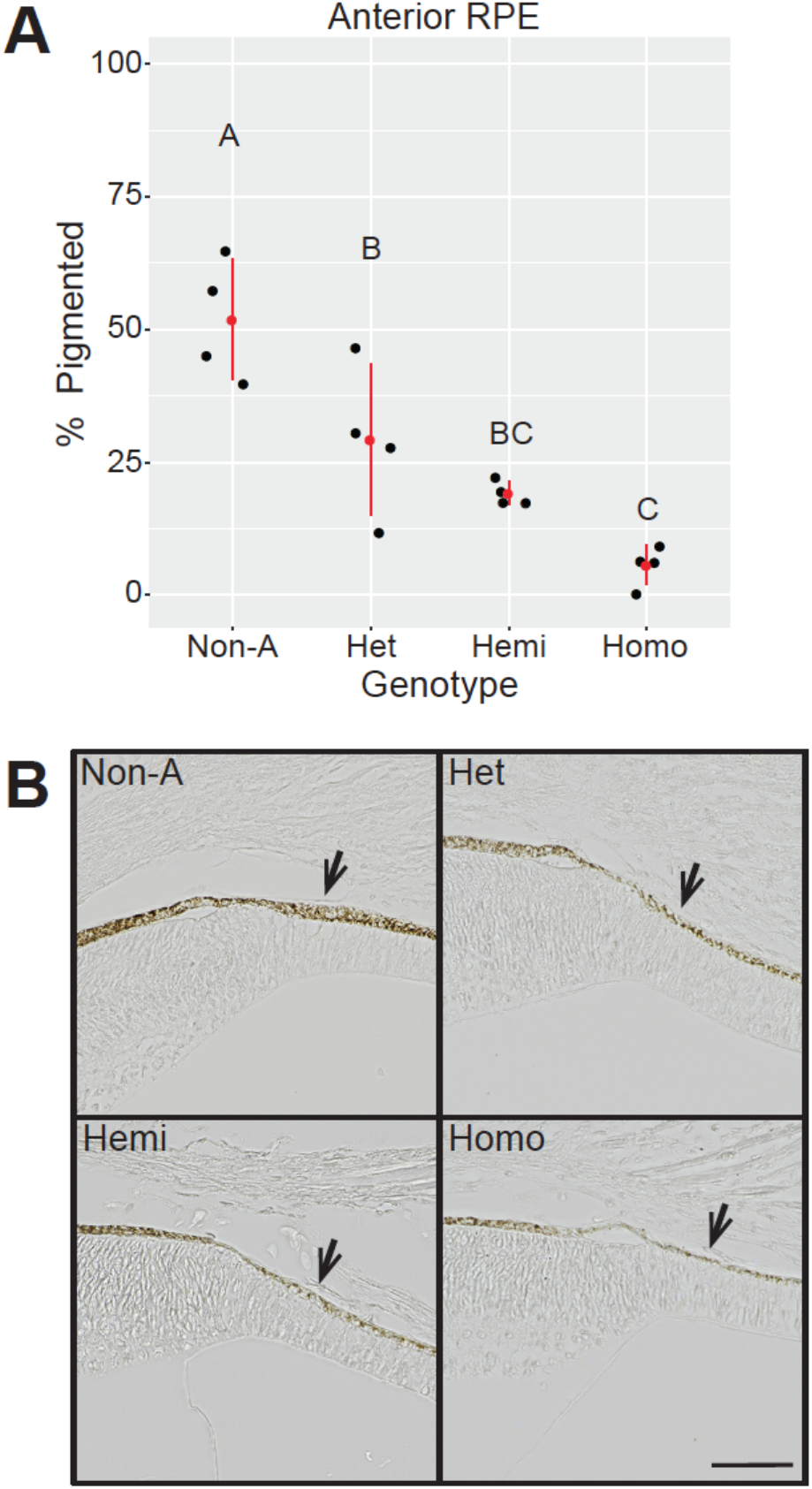
Anterior RPE pigmentation is reduced in all Almond genotypes. (A) Percentage of the anterior RPE that is pigmented in different genotypes. Four hatchling eyes were analyzed per genotype. Three sections were analyzed for each eye, and mean values are plotted as black dots. Red dot and line show mean and standard deviation, respectively. Different letters indicate groups with statistically significant differences in gene expression determined by Anova and post-hoc Tukey (p < 0.05). Genotypes: Non-A, non-Almond; Het, heterozygous Almond; Hemi, hemizygous Almond; Homo, homozygous Almond. (B) Representative images of the RPE for each genotype. Arrows point to the pigmented layer of the RPE. Pigmentation is reduced in all Almond genotypes. Scale bar in bottom right image is 50 μm.

### Posterior RPE

The posterior RPE (Fig 1D) adjacent to the optic nerve – the most posterior eye structure measured – had smaller pigmentation differences among genotypes than other regions. Non-Almond, heterozygous Almond, and hemizygous Almond genotypes were similar to each other with mean pigmented areas of 66.0%, 53.3%, and 56.2%, respectively (post-hoc Tukey, p > 0.05; Fig 5A). Only homozygous Almonds had significantly less pigment than non-Almonds with a mean pigmented area of 35.9 % (Fig 5A and 5B; Anova, p = 0.01; post-hoc Tukey, p = 0.0064).

**Figure 5.**
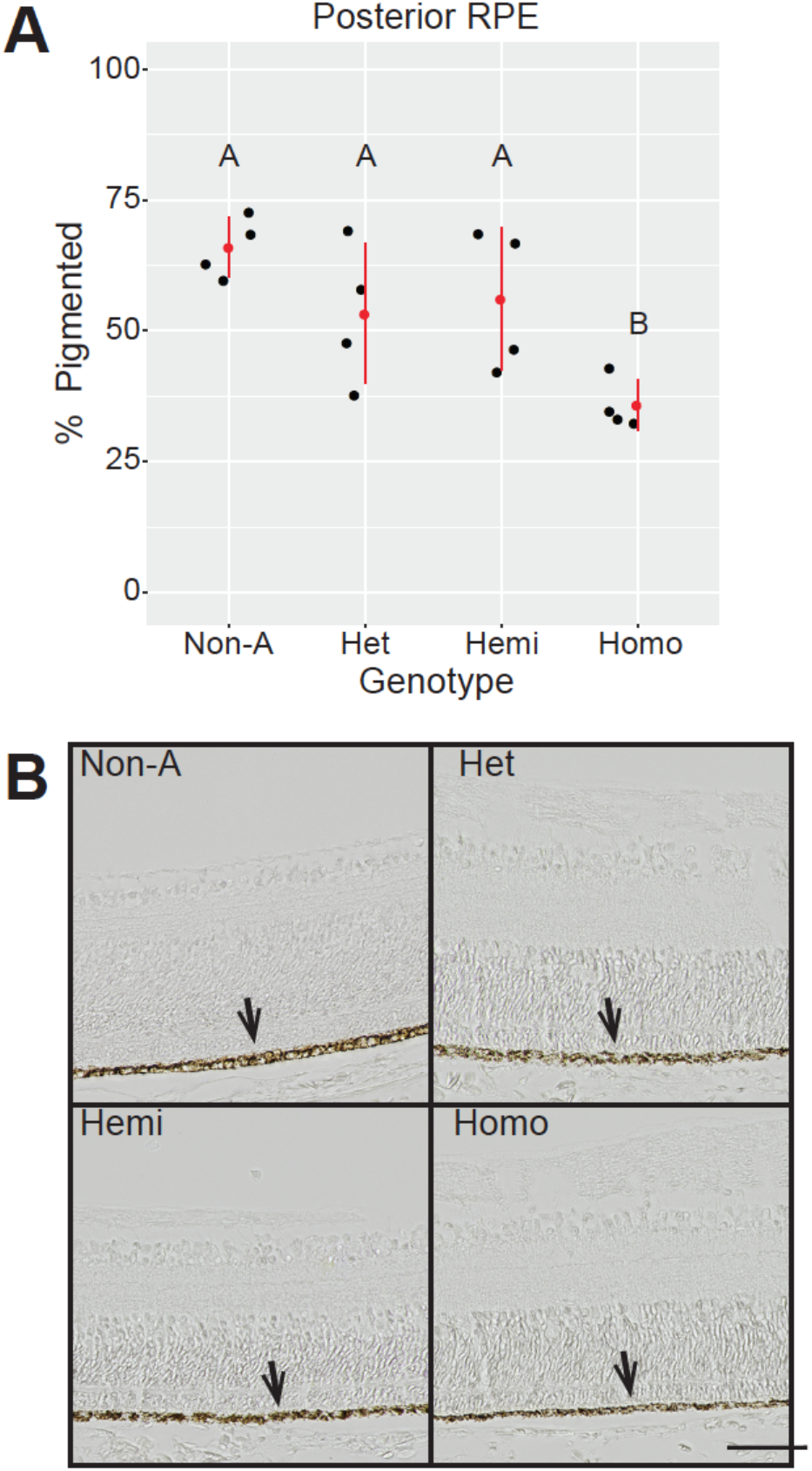
Posterior RPE pigmentation is less affected than anterior pigmentation in eyes of Almond pigeons. (A) Percentage of the RPE that is pigmented in different genotypes. Four hatchling eyes were analyzed per genotype. 2-3 sections were analyzed for each eye, and mean values are plotted as black dots. Red dot and line show mean and standard deviation, respectively. Different letters indicate groups with statistically significant differences in gene expression determined by Anova and post-hoc Tukey (p < 0.05). Genotypes: Non-A, non-Almond; Het, heterozygous Almond; Hemi, hemizygous Almond; Homo, homozygous Almond. (B) Representative images of the RPE for each of genotype. Arrows point to the pigmented layer of the RPE. Pigmentation is only slightly reduced in heterozygous Almond hatchling eyes. Scale bar in bottom right image is 50 μm.

### Pigment-containing cell layers

By gross visual examination, all histologic sections of all genotypes appeared to retain the epithelial layer that contains pigment in non-Almond birds. Even in homozygous Almond hatchlings, which showed the least amount of pigmentation in all eye regions, this layer also did not have any obvious morphological changes compared to non-Almond.

In summary, all of the Almond genotypes had significant decreases in pigmentation in anterior structures, especially in the iris and ciliary body, the two most anterior structures. In contrast, pigmentation in the posterior RPE was significantly lower only in homozygous Almond eyes.

## Discussion

### Almond alleles cause more anterior than posterior depigmentation

In the eyes of hatchling pigeons with one or two Almond alleles, anterior structures had greater reductions in pigmentation than did posterior structures. A potential explanation for these disparities is the different developmental origins of melanin-depositing cells in different segments of the eye. In the posterior RPE, pigment is deposited by cells of the optic cup neuroepithelium [20]. In anterior eye structures, including the iris and ciliary body, pigment comes from both the neural crest-derived melanocytes and cells that originate from the optic cup neuroepithelium [7]. In birds with Almond alleles, we found that structures receiving pigment from both sources were less pigmented than those receiving pigment from the neuroepithelium alone. Therefore, neural crest-derived pigmentation of the eye is more severely affected than neuroepithelium-derived pigmentation. Similarly, feather pigmentation is deposited by neural crest-derived melanocytes, and plumage color is severely affected in birds with Almond alleles. Thus, Almond-linked pigmentation changes in the anterior segment of the eye and the epidermis share a common cellular origin, the neural crest.

In the posterior RPE, we saw a significant but relatively modest pigmentation reduction in homozygous Almond eyes relative to all other genotypes. This situation is analogous to what is observed in the plumage: hemizygous and heterozygous Almond pigeons have reduced (but still plentiful) feather pigmentation, but homozygotes lose almost all plumage pigmentation [36]. In developing homozygous feathers, but not in other genotypes, we saw a collapse of the pigment synthesis gene expression pathway. We do not have comparable gene expression data for eye structures, but we speculate that a similarly broad set of pigment synthesis changes impact all pigment-producing cells, not just those derived from the neural crest. Overall, in both the feathers and eyes of Almond pigeons, we see a dosage effect wherein two Almond alleles cause a more severe pigmentation phenotype than one. For reasons that we do not yet understand, anterior eye pigment deposition is more sensitive to this dosage effect than posterior deposition. Melanocytes are migratory and encounter dynamic cellular surroundings as they move toward their destination, whereas RPE cells have a more stable tissue microenvironment; perhaps these contextual differences contribute to differential resiliency of pigment cell development and function [7].

### Homozygous Almond eyes have both pigmentation and structural defects

Homozygous Almond eyes had the least pigmentation of any genotype. In the most anterior regions, the iris and ciliary body, almost no pigment was visible (Fig 2B and 3B). Homozygous Almond was also the only genotype with significantly reduced pigmentation in the posterior RPE (Fig 5A and 5B). It remains unclear if and how a paucity of melanin is mechanistically linked to other structural developmental defects such as bulging “pop-eyes”, colobomas, and the absence of lenses [31–35]. Neural crest cells are the progenitors of melanocytes and other structures that comprise the anterior segment of the eye [7, 56–58]. Hence, if the Almond mutation specifically affects neural crest-derived structures, then it could lead to the anterior segment defects observed in homozygous Almond pigeons. A related possibility is that pigment deposition could be necessary for the normal development of anterior segment structures. Signaling between tissue layers of the eye is necessary for proper development [59]. Therefore, defects in anterior pigmented structures could, in turn, lead to bulging or malformed corneas, as well as lens and eyelid defects.

Another potential route to eye structural defects is the “leaking” of toxic pigment cell components. In humans, pigmentary glaucoma (PG) and pigment dispersal syndrome (PDS) affect anterior eye structures and are sometimes associated with mutations in *PMEL* [60], PMEL is critical for melanosome maturation and physically interacts with MLANA [38], a protein that is encoded in the Almond genomic candidate region of pigeons [36]. In cases of PDS, pigment disperses from the iris to surrounding tissues and induces other defects, such as increased ocular pressure and corneal endothelial pigment deposition [61]. However, both PDS and PG generally do not affect lens development, which precedes pigment deposition.

### Lessons from melanosome defects in other animals

Links between epidermal pigmentation changes and eye defects have been observed in several animal species. In some cases, the genes responsible for these linked defects are involved in the function or structure of the melanosome, including *Slc45a2, Rab38*, and *Pmel* [1, 3, 40, 62–64], Of particular interest, mutations in *Pmel* are associated with changes in both epidermal pigmentation and eye or vision defects in several species. *Pmel* encodes an amyloid structural protein that contributes to the melanosome matrix [65, 66]. Pmel interacts directly with Mlana, a top candidate gene for Almond in pigeons, during melanosome maturation [38].

*Pmel* mutant animals have varying degrees of epidermal pigmentation changes and eye defects, and their phenotypes are often qualitatively similar to Almond pigeons. In zebrafish, loss of function in one *pmel* paralog (*pmela*) leads to a reduction in pigment in the body and eye, and also enlarged anterior eye segments, likely due to increased intraocular pressure [60]. Similarly, some homozygous Almond pigeons also appear to have enlarged anterior segments [30, 69, 70, personal observations]. Other *Pmel* mutants, including silver horses [48, 68, 69] and homozygous Merle dogs [16, 17, 43, 70], also have associated defects in anterior eye structures. A counterexample of this trend is the *PMEL17* Dominant white mutation in chickens, which results in the loss of epidermal feather pigmentation but is not associated with changes in eye pigmentation [25].

Based on our findings of an anterior-posterior pigmentation gradient in Almond pigeon eyes, we are interested to test whether eye pigmentation disparities in other species track with embryological origins of pigment cells. For example, we predict that *pmel* mutants with epidermal and anterior eye structural defects, such as Merle dogs and silver horses, might have anterior-posterior gradients of eye pigmentation defects that are similar to what we observe in Almond pigeons.

## Materials and Methods

### Ethics statement

Animal husbandry and experimental procedures were performed in accordance with protocols approved by the University of Utah Institutional Animal Care and Use Committee (protocols 10-05007, 13-04012, and 16-03010).

### Sample collection

Offspring from crosses of heterozygous Almond males (Z^*St*^/Z^+^) and Almond females (Z^*St*^/W) were euthanized within 8 hr of hatching. Whole eyes were collected and fixed overnight in 4% paraformaldehyde, then preserved in 70% ethanol. Eye samples for 4 individuals of each genotype (non-Almond, hemizygous Almond, heterozygous Almond, and homozygous Almond) were then embedded in paraffin and sagittally sectioned to a thickness of 8 μm by ARUP Laboratories (Salt Lake City, UT).

### Genotyping of Hatchlings

Brain tissue was collected at the same time as each eye sample and stored at −80°C for DNA extraction and genotyping. We extracted DNA using the Qiagen DNeasy Blood & Tissue Kit, following the manufacturer’s protocol (Qiagen Sciences, Germantown, MD). Sex was determined using a PCR assay [71]. Copy number at the Almond locus was estimated using a custom TaqMan Copy Number Assay (ThermoFisher Scientific, Waltham, MA) targeted to an intron in *Mlana* (MLANA_CCWR201), and an intron in *RNaseP* was used for normalization. TaqMan samples were run in quadruplicate according to manufacturer’s protocol. Copy number (CN) was determined using CopyCaller Software v2.1 (ThermoFisher Scientific, Waltham, MA). Two separate plates of DNA samples were included in the TaqMan assay. The first plate contained all but one sample used in this study. We also included DNA control samples from adults that were confirmed by phenotype and genotype to be non-Almond (3 samples), heterozygous Almond (2 samples), and hemizygous Almond (2 samples). Non-Almond genotypes were called in our experimental samples when the calculated CN was less than 2.0. Heterozygous and hemizygous Almond genotypes were called when CN was between 2.0 and 12.0, and the distinction between these genotypes was based on a PCR assay for sex (hemizygous Almond if female, heterozygous Almond if male). The second assay plate contained only one experimental sample, KD710 NOV17#04, plus Almond and non-Almond controls (one of each). This plate yielded lower calculated CN results for heterozygous Almond individuals, and thus genotype prediction results were adjusted accordingly (indicated with * in S1 Table). Results for sex and copy number genotyping for all samples from both plates are shown in S1 Table.

### Imaging of eye sections

We did not obtain clear anterior and posterior landmarks together in the same histologic sections, so we used different sections to quantify anterior and posterior pigmentation for each eye. For pigmentation analyses of anterior structures, we selected 3 sections from each eye that included the approximate center of the lens. Images were taken of the iris pigmented epithelium, the ciliary body, and the ora serrata (for anterior RPE analysis). For pigmentation analyses of the posterior RPE, we selected 2-3 sections containing the optic nerve, and images were taken of the RPE on both sides adjacent to the optic nerve. Sections were imaged using a Nikon Ti-E inverted microscope (Nikon Instruments Inc, Melville, NY) with a high-sensitivity Andor Zyla sCMOS camera (Oxford Instruments, Abingdon, Oxon, United Kingdom) and LED light source. For the IPE, anterior RPE, and posterior RPE images were taken using 100x magnification, and Z-stacks of each section were taken using 0.02-μm steps through the tissue layers containing pigmented melanosomes using Nikon NIS-Elements software. For the ciliary body, images were taken on the same microscope using 20x magnification and 0.2 μm steps. Stacks were merged to create an Extended Depth of Focus (EDF) file. EDF files were then exported to create a TIFF file for image analysis.

### Image analysis

TIFF images were imported into Adobe Photoshop (Adobe Inc., San Jose, California). For each image, the IPE or RPE structure was selected and the background (non-IPE or non-RPE tissues) was deleted. For the ciliary body, area for analysis was determined by cropping regions that contained pigmented melanosomes or reflective structures that are smaller than a single cell, which include melanosomes and pre-melanosomes. The cropped images were imported into ImageJ (imagej.nih.gov/) and the threshold command was used to determine %-pigmented area for each structure. Optimal thresholding values for pigmented area were determined visually in ImageJ using non-Almond samples for reference, to find values that include the most melanosomes possible, without picking up background pixels (0-100 for the 100x images of IPE or RPE layer, and 0-120 for the 20x images of the ciliary body). Thresholding values for total area were determined visually to include the total area of the cropped regions (0-220 for the 100x images of the IPE and RPE, and 0-240 for the 20x images of the ciliary body). The %-pigmented area for each structure was determined by dividing the pigmented area by the total area. See S2 Table for raw data.

## Acknowledgements

We thank the Cell Imaging Core at the University of Utah for use of microscopy equipment and software; Michael J. Bridge for assistance and advice on imaging; Kristen Kwan for discussions of eye development; Anna Vickrey for assistance with thresholding analysis and comments on the manuscript; and Emily Maclary and Elena Boer for comments on the manuscript. This work was supported by NSF fellowship GRF 1256065 to R.B. and NIH grant R35 GM131787 to M.D.S. The funders had no role in study design, data collection and analysis, decision to publish, or preparation of the manuscript.

